# A Large-Scale ENIGMA Multisite Replication Study of Brain Age in Depression

**DOI:** 10.1101/2022.08.29.505635

**Authors:** Laura K.M. Han, Richard Dinga, Ramona Leenings, Tim Hahn, James H. Cole, Lyubomir Aftanas, Alyssa R. Amod, Bianca Besteher, Romain Colle, Emmanuelle Corruble, Baptiste Couvy-Duchesne, Konstantin Danilenko, Paola Fuentes-Claramonte, Ali Saffet Gonul, Ian H. Gotlib, Roberto Goya-Maldonado, Nynke A. Groenewold, Paul Hamilton, Naho Ichikawa, Jonathan C. Ipser, Eri Itai, Sheri-Michelle Koopowitz, Meng Li, Go Okada, Yasumasa Okamoto, Churikova Olga, Evgeny Osipov, Brenda W.J.H. Penninx, Edith Pomarol-Clotet, Elena Rodríguez-Cano, Matthew D. Sacchet, Hotaka Shinzato, Kang Sim, Dan J. Stein, Aslihan Uyar-Demir, Dick J. Veltman, Lianne Schmaal

## Abstract

**Background:** Several studies have evaluated whether depressed persons have older appearing brains than their nondepressed peers. However, the estimated neuroimaging-derived “brain age gap” has varied from study to study, likely driven by differences in training and testing sample (size), age range, and used modality/features. To validate our previously developed ENIGMA brain age model and the identified brain age gap, we aim to replicate the presence and effect size estimate previously found in the largest study in depression to date (N=2,126 controls & N=2,675 cases; +1.08 years [SE 0.22], Cohen’s d=0.14, 95% CI: 0.08-0.20), in independent cohorts that were not part of the original study.

**Methods:** A previously trained brain age model (www.photon-ai.com/enigma_brainage) based on 77 FreeSurfer brain regions of interest was used to obtain unbiased brain age predictions in 751 controls and 766 persons with depression (18-75 years) from 13 new cohorts collected from 20 different scanners.

**Results:** Our ENIGMA MDD brain age model generalized reasonably well to controls from the new cohorts (predicted age vs. age: *r* = 0.73, *R*^2^=0.47, MAE=7.50 years), although the performance varied from cohort to cohort. In these new cohorts, on average, depressed persons showed a significantly higher brain age gap of +1 year (SE 0.35) (Cohen’s d□=□□.15, 95% CI: 0.05–0.25) compared with controls, highly similar to our previous finding.

**Conclusions:** This study further validates our previously developed ENIGMA brain age algorithm. Importantly, we replicated the brain age gap in depression with a comparable effect size. Thus, two large-scale independent mega-analyses across in total 32 cohorts and >3,400 patients and >2,800 controls worldwide show reliable but subtle effects of brain aging in adult depression.

## 1. INTRODUCTION

Recently, considerable literature has emerged around the theme of human aging. Aging is accompanied by complex biological changes, such as linear and nonlinear brain structural changes (Anderton, 2002). Machine learning algorithms can leverage these age-related brain patterns to predict **chronological age**, to explain individual differences in aging. If (structural) magnetic resonance imaging (MRI) data are used as input for these algorithms, the output can be considered as an estimate of brain-based biological age, or, predicted **brain age** (Cole and Franke, 2017). Over the past decade there has been an exponential increase in studies investigating brain age (Baecker et al., 2021), with this metric being used to quantify one’s brain health state, as well as risk for aging-related diseases and mortality (Cole et al., 2018). These are important indicators of neurodegenerative diseases such as Alzheimer’s or multiple sclerosis; however, these risks are also commonly increased (albeit to a lesser extent) in major depressive disorder (MDD) (Penninx, 2017).

The estimated neuroimaging-derived “brain age gap” (predicted brain age minus chronological age, i.e., brain-predicted age difference, or, **brain-PAD**) in depression varies across studies in terms of both effect size and statistical significance. These differences are likely driven by differences in sample properties (e.g., age range), but also training and testing sample (size), and methods used (e.g., modality/features). A recent systematic review and meta-analysis summarized that the majority (4 out of 7) of the existing studies of brain-PAD in depression did not establish a significant case-control difference (Ballester et al., 2022). Yet, effects were compatible across studies; thus, all studies identified a higher average brain age gap in depression compared to controls, with a pooled effect of approximately +1 year of added aging, although estimated gaps ranged from 0.13 to 4.92 years. Our previous ENIGMA MDD consortium study, the largest study of brain age in depression to date (N=2,126 controls and N=2,675 patients), showed a +1.08 year higher brain-PAD in depression (Cohen’s d=0.14), but with no evidence that this effect was driven by specific (clinical) characteristics such as age, age of onset, recurrence status, remission status, or antidepressant use (Han et al., 2021a). The subsequent addition of new cohorts to the ENIGMA MDD consortium since our previous brain age study provides us with an unique opportunity to perform an independent replication study in new data to validate our developed algorithm, as well as determine whether depression is consistently associated with older appearing brains (Wrigglesworth et al., 2021). Studying the impact of depressive psychopathology on age-related structural brain patterns may help to explain why persons with depression have an increased risk for poorer brain and physical health compared to their nondepressed peers.

Our ENIGMA brain age algorithm was trained on 952 male and 1,236 female healthy controls from 19 cohorts. We used 77 FreeSurfer-derived ROI features (34 cortical thickness, 34 cortical surface area, 7 subcortical volumes, lateral ventricles, and intracranial volume) to predict chronological age using ridge regression. While other existing (deep neural network) algorithms may potentially provide more accurate predictions (Lombardi et al., 2020), most of them rely on using higher-dimensional imaging data as input (e.g., raw scans, individual-level voxels, or vertices). Within the ENIGMA consortium, many cohorts have shared data in the form of brain-derived summary measures (i.e., FreeSurfer brain regions of interest, or ROIs). The current FreeSurfer ROIs method is thus one of the more practical ways to perform a large multisite brain age mega-analysis in depression, facilitated by the collaborative nature of the ENIGMA MDD working group. Our first study showed good out-of-sample generalization to new and unseen controls and patients from the same cohorts as the model was trained on, as well as completely independent controls from cohorts not included in training (i.e., ENIGMA Bipolar Disorder controls)(Han et al., 2021a). Additionally, other ENIGMA studies have further demonstrated the validity of this model (Clausen et al., 2022), identifying a higher brain-PAD in schizophrenia (Constantinides et al., 2022).

This study aims to further validate the ENIGMA FreeSurfer ROI-based brain age prediction method, by evaluating the performance of our algorithm in 13 new and unseen cohorts of individuals with depression collected from 20 independent scanners. Importantly, we aim to contribute to the growing area of brain age research by attempting to replicate the magnitude of the brain age gap difference previously reported by the ENIGMA-MDD consortium between persons with depression (N=766) and controls (N=751) using this method.

## 2. METHODS

### Samples

Thirteen independent cohorts (N=1,517) from the ENIGMA MDD working group with data from people with major depression and controls (18-75 years old) participated in this replication study. Cohort-specific details on demographics, basic clinical characteristics, and exclusion criteria can be found in the **Supplement**. All sites obtained approval from their local institutional review boards and ethics committees. All study participants provided written informed consent.

### ENIGMA brain age prediction model

Model development is described in more detail in (Han et al., 2021a), but in short, ridge regression was used to predict age from 77 FreeSurfer-derived features (7 subcortical volumes, 34 cortical thickness regions, 34 cortical surface area regions, lateral ventricles and intracranial volume) in healthy controls (no history of mental or neurological illness). Separate models were trained for male (N=952) and female (N=1,236) controls. The ENIGMA brain age model is publicly available (www.photon-ai.com/enigma_brainage) and was applied to the independent new ENIGMA MDD cohorts included in the current study. A schematic overview is displayed in **Figure 1.**

**Figure 1.**
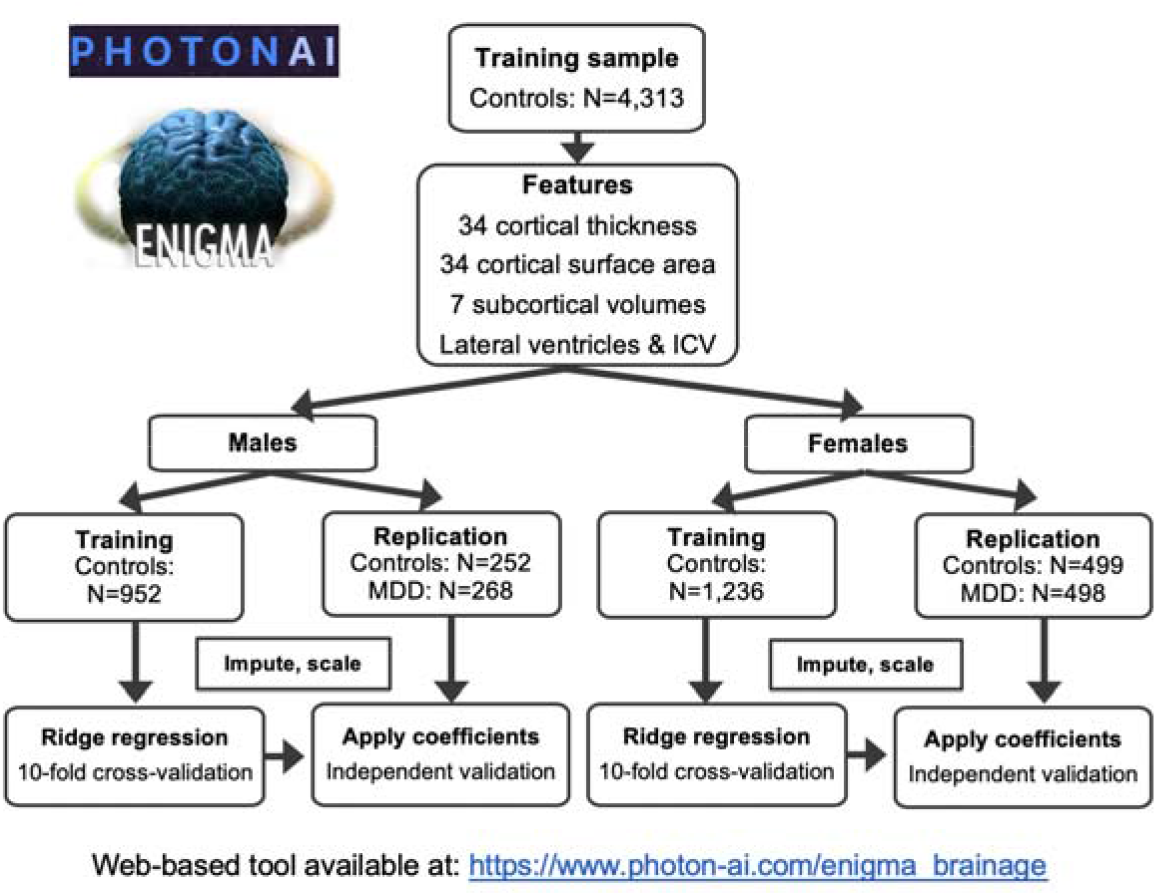
Schematic overview of features used and data used to train the ENIGMA brain age model. Learned sex-specific ridge regression coefficients were applied to the current independent replication test data.

### Model generalization

The ENIGMA brain age prediction model has previously been validated in 646 unseen male and 757 unseen female control samples from 23 independent scanners that were not part of the training data (Han et al., 2021a), as well as in other disease working groups of ENIGMA (Clausen et al., 2022; Constantinides et al., 2022). In the current study, model generalization was evaluated in control samples collected from 20 additional independent scanners (N=252 males and N=499 females). To assess model performance in these data acquired from completely independent cohorts, we calculated *(1)* mean absolute error (MAE), *(2)* weighted MAE (i.e., _w_MAE, an age range informed metric;), *(3)* Pearson correlation coefficients between predicted brain age and chronological age, and *(4)* the proportion of the variance explained by the model (*R^2^*). *R^2^* was calculated using the caret package according to the formula:

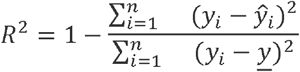

Where *n* is the number of subjects, y is the chronological age, 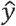 is the predicted age, and *<u>y</u>* is the average age of subjects in the test set. Please note that according to this formula, *R*^2^ can be negative, even if a correlation between age and predicted age is positive. This happens when model predictions have larger errors than predicting the average age, such as when predictions are biased or have a relatively large variance.

### Statistical analyses

A mega-analytic approach was taken to replicate the presence and size of the brain age gap in depression by pooling data across all thirteen new cohorts. The brain age gap (predicted brain-based age minus chronological age, or, **brain-PAD**) was calculated for each individual and used as the outcome variable in analyses comparing the difference between brain-PAD in people with depression and controls and examining associations between brain-PAD and clinical characteristics. Each dependent measure of the i^th^ individual at j^th^ scanning site was modeled as follows:

1. Brain-PAD_ij_□=□intercept□+□β_1_(Dx)□+□β_2_(sex)□+□β_3_(age)□+□β_4_(age^2^)□+□β_5_(Dx□×□age)□+□β_6_(Dx□×□sex)□+□β_7_(age□×□sex)□+□β_8_(Dx□×□age□×□sex)□+□U_j_□+□ε_ij_
2. Brain-PAD_ij_□=□intercept□+□β_1_(Dx)□+□β_2_(sex)□+□β_3_(age)□+□β_4_(age^2^)□+□β_5_(Dx□×age)□+□β_6_(Dx□×sex) + U_j_−__+□ε_ij_
3. Brain-PAD_ij_□=□intercept□+□β_1_(Dx)□+□β_2_(sex)□+□β_3_(age)□+□β_4_(age^2^)□+□U_j_□+□ε_ij_

Intercept, Dx (MDD diagnosis), sex, and all age effects were fixed. The terms Uj and ε_ij_ are normally distributed and represent the random intercept attributed to the scanning site and the residual error, respectively. Standardized Cohen’s *d* was calculated to indicate the size of the effect. Within the patient group, we also used linear mixed models to examine brain-PAD associations with clinical characteristics (i.e., recurrence status [first versus recurrent episode], antidepressant (AD) status [taking AD yes/no at time of scanning], remitted status [acute versus remitted], age of onset of depression [categorized as: early, <26 years; middle adulthood, >25 and <56 years; and late adulthood onset, >55 years]). In addition to the **mega**-analytic approach, a **meta**-analytic approach was also performed to provide further insights into the generalization of the ENIGMA brain age model and case-control difference in individual cohorts. Exploratory effects of cohort specific characteristics (i.e., sample size, mean age, proportion of females) but also potential (scan) technical moderators (i.e., FreeSurfer version, field strength, scanner vendor, performance accuracy metrics [MAE, R^2^]) on the brain-PAD outcome were examined by random effects meta-regressions analyses using the *metafor* package in *R* (a more detailed description on methods can be found in the **Supplementary Methods**). Replication analyses were tested one-sided, whereas analyses within the patient group and meta-regressions were tested two-sided. All statistical tests were considered significant at *p*<0.05.

## 3. RESULTS

### 3.1. Participants

Participant characteristics are presented in **Table 1**. Thirty individuals from one cohort were excluded from the study based on having >10% missing structural brain ROIs data. Two individuals >75 years old from another cohort were also excluded from this study, given that the model was trained on data restricted within 18-75 years. Eight persons showed a brain-PAD with a calculated Z-score >3 (i.e., >3SD away from the global mean) and were excluded from analysis. In total, we included data from N=1,517 participants, including N=751 controls (66% females) and N=766 persons with (current) MDD (65% females). The **Supplement** includes cohort-specific information on participants (**Table S1**), image acquisition and processing (**Table S2**) and instruments used for depression ascertainment (**Table S3**).

**Table 1.** Participant Characteristics per Diagnostic Group.

| Characteristic |  | N | Controls, N = 751<sup>1</sup> | MDD, N = 766<sup>1</sup> |
| --- | --- | --- | --- | --- |
| *Cohort* | AFFDIS | 49 | 20 (2.7%) | 29 (3.8%) |
|  | CSAN | 107 | 49 (6.5%) | 58 (7.6%) |
|  | DCHS | 70 | 54 (7.2%) | 16 (2.1%) |
|  | FIDMAG | 69 | 34 (4.5%) | 35 (4.6%) |
|  | Hiroshima | 315 | 167 (22%) | 148 (19%) |
|  | TiPS | 105 | 75 (10.0%) | 30 (3.9%) |
|  | MOODS | 96 | 32 (4.3%) | 64 (8.4%) |
|  | MOTAR | 108 | 68 (9.1%) | 40 (5.2%) |
|  | NESDA | 219 | 65 (8.7%) | 154 (20%) |
|  | Novo | 128 | 52 (6.9%) | 76 (9.9%) |
|  | Singapore | 40 | 17 (2.3%) | 23 (3.0%) |
|  | SoCAT | 179 | 100 (13%) | 79 (10%) |
|  | StanfFAA | 32 | 18 (2.4%) | 14 (1.8%) |
| Chronological age (years) |  | 1,517 | 38.80 ± 12.89 (17.00-73.00) | 39.82 ± 12.70 (18.00-73.00) |
| Predicted brain age (years) |  | 1,517 | 41.75 ± 10.79 (14.78-74.60) | 43.94 ± 10.65 (15.22-75.66) |
| Brain-PAD (years) |  | 1,517 | 2.95 ± 8.89 (-25.31-31.39) | 4.12 ± 9.86 (-25.19-32.40) |
| Sex | Female | 1,517 | 499 (66%) | 498 (65%) |
| *Recurrent status* | First | 1,130 |  | 225 (38%) |
|  | Recurrent |  |  | 375 (62%) |
| *Antidepressant use* | AD-free | 1,131 |  | 357 (59%) |
|  | AD-using |  |  | 244 (41%) |
| *Remitted status* | Remitted | 1,004 |  | 41 (7.5%) |
|  | Acute |  |  | 508 (93%) |
| *FreeSurfer version* | 5 | 1,517 | 468 (62%) | 490 (64%) |
|  | 6 |  | 134 (18%) | 139 (18%) |
|  | 7 |  | 149 (20%) | 137 (18%) |
| *Field strength* | 1.5T | 1,517 | 34 (4.5%) | 35 (4.6%) |
|  | 3T |  | 717 (95%) | 731 (95%) |
| *Scanner vendor* | GE | 1,517 | 104 (14%) | 125 (16%) |
|  | Philips |  | 182 (24%) | 281 (37%) |
|  | Siemens |  | 465 (62%) | 360 (47%) |
| <sup>1</sup> n (%); Mean ± SD (Minimum-Maximum) |  |  |  |  |

### 3.2. Brain age prediction performance

Model generalizability was evaluated in the healthy control samples. Both pooled and cohort-specific model performances are presented in **Table 2.** While the generalization power varied from cohort to cohort, the pooled performance accuracy was comparable to the out-of-sample generalizability previously reported in (Han et al., 2021a), with current metrics between predicted brain age and chronological age of Pearson’s *r*=0.73, *R^2^*=.47, MAE=7.50 years, and _w_MAE=0.13. **Figure 2** shows the predicted brain age against chronological age per cohort (separate regression lines for controls and patients). Cohorts showing a negative *R^2^* showed negative mean cortical thickness deviations compared to the grand mean of combined cohorts (**Supplementary Figure S1**). Performance metrics in patient samples can also be found in the **Supplementary Table S4.**

**Table 2.**
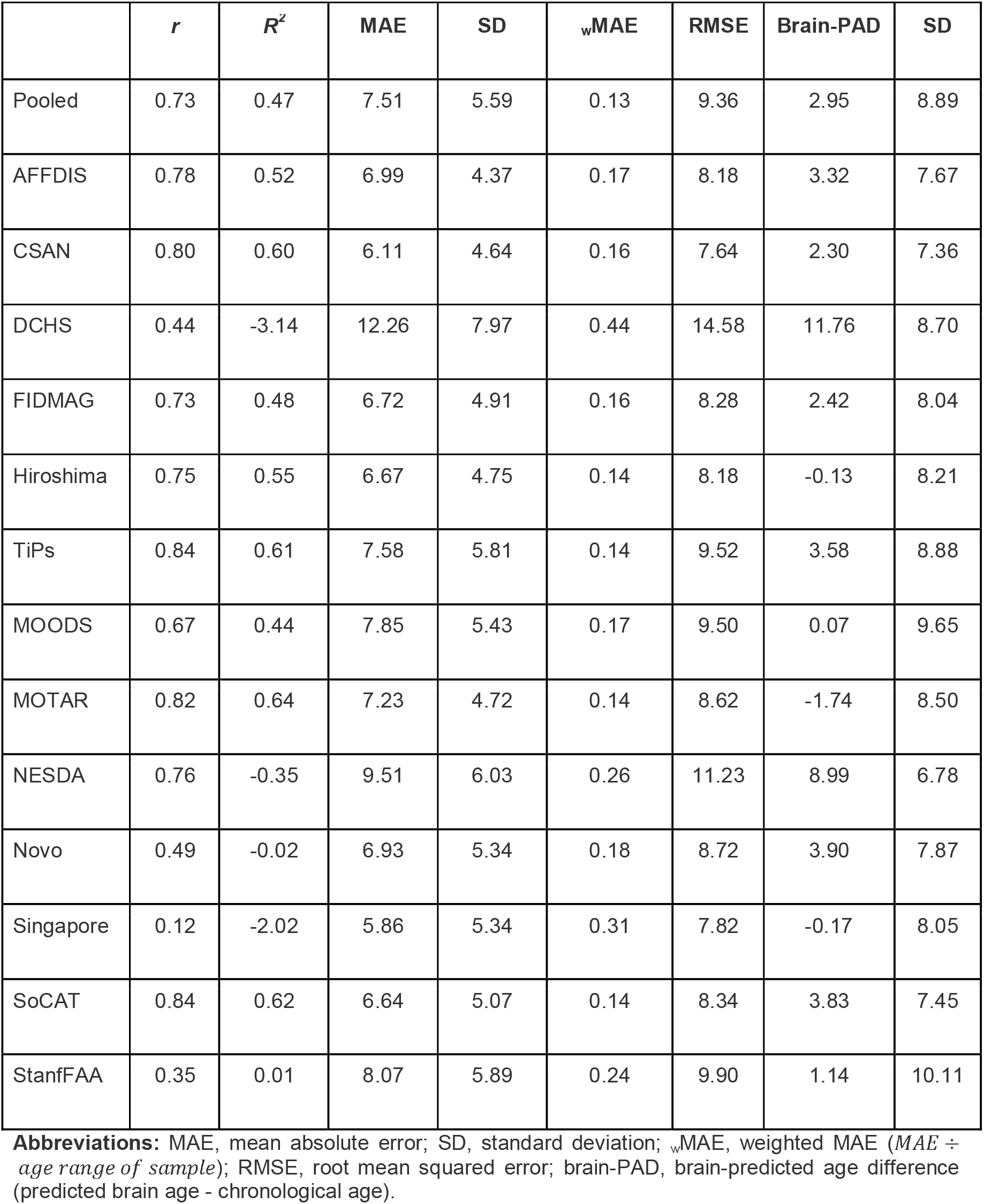
Generalizability to independent control cohorts.

|  | *r* | *R*<sup>2</sup> | MAE | SD | <sub>w</sub>MAE | RMSE | Brain-PAD | SD |
| --- | --- | --- | --- | --- | --- | --- | --- | --- |
| Pooled | 0.73 | 0.47 | 7.51 | 5.59 | 0.13 | 9.36 | 2.95 | 8.89 |
| AFFDIS | 0.78 | 0.52 | 6.99 | 4.37 | 0.17 | 8.18 | 3.32 | 7.67 |
| CSAN | 0.80 | 0.60 | 6.11 | 4.64 | 0.16 | 7.64 | 2.30 | 7.36 |
| DCHS | 0.44 | -3.14 | 12.26 | 7.97 | 0.44 | 14.58 | 11.76 | 8.70 |
| FIDMAG | 0.73 | 0.48 | 6.72 | 4.91 | 0.16 | 8.28 | 2.42 | 8.04 |
| Hiroshima | 0.75 | 0.55 | 6.67 | 4.75 | 0.14 | 8.18 | -0.13 | 8.21 |
| TiPs | 0.84 | 0.61 | 7.58 | 5.81 | 0.14 | 9.52 | 3.58 | 8.88 |
| MOODS | 0.67 | 0.44 | 7.85 | 5.43 | 0.17 | 9.50 | 0.07 | 9.65 |
| MOTAR | 0.82 | 0.64 | 7.23 | 4.72 | 0.14 | 8.62 | -1.74 | 8.50 |
| NESDA | 0.76 | -0.35 | 9.51 | 6.03 | 0.26 | 11.23 | 8.99 | 6.78 |
| Novo | 0.49 | -0.02 | 6.93 | 5.34 | 0.18 | 8.72 | 3.90 | 7.87 |
| Singapore | 0.12 | -2.02 | 5.86 | 5.34 | 0.31 | 7.82 | -0.17 | 8.05 |
| SoCAT | 0.84 | 0.62 | 6.64 | 5.07 | 0.14 | 8.34 | 3.83 | 7.45 |
| StanfFAA | 0.35 | 0.01 | 8.07 | 5.89 | 0.24 | 9.90 | 1.14 | 10.11 |

**Abbreviations:** MAE, mean absolute error; SD, standard deviation; <sub>w</sub>MAE, weighted MAE (*MAE ÷ age range of sample*); RMSE, root mean squared error; brain-PAD, brain-predicted age difference (predicted brain age - chronological age).

**Table 3.**
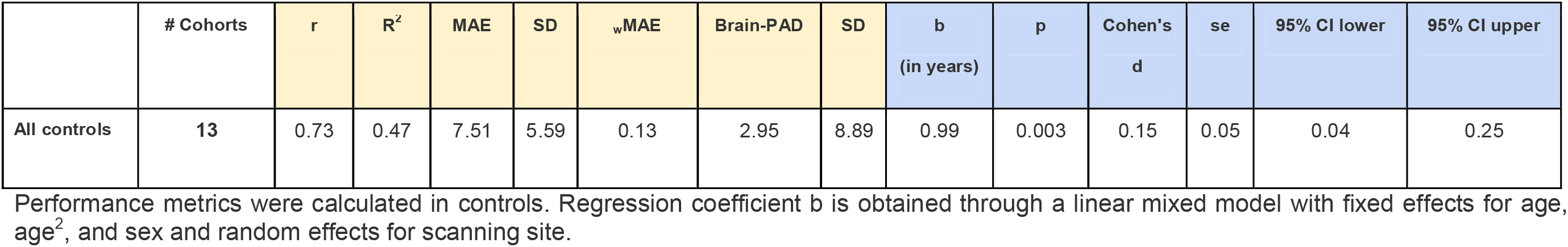
Pooled performance accuracy and case-control difference in brain-PAD.

**Figure 2.**
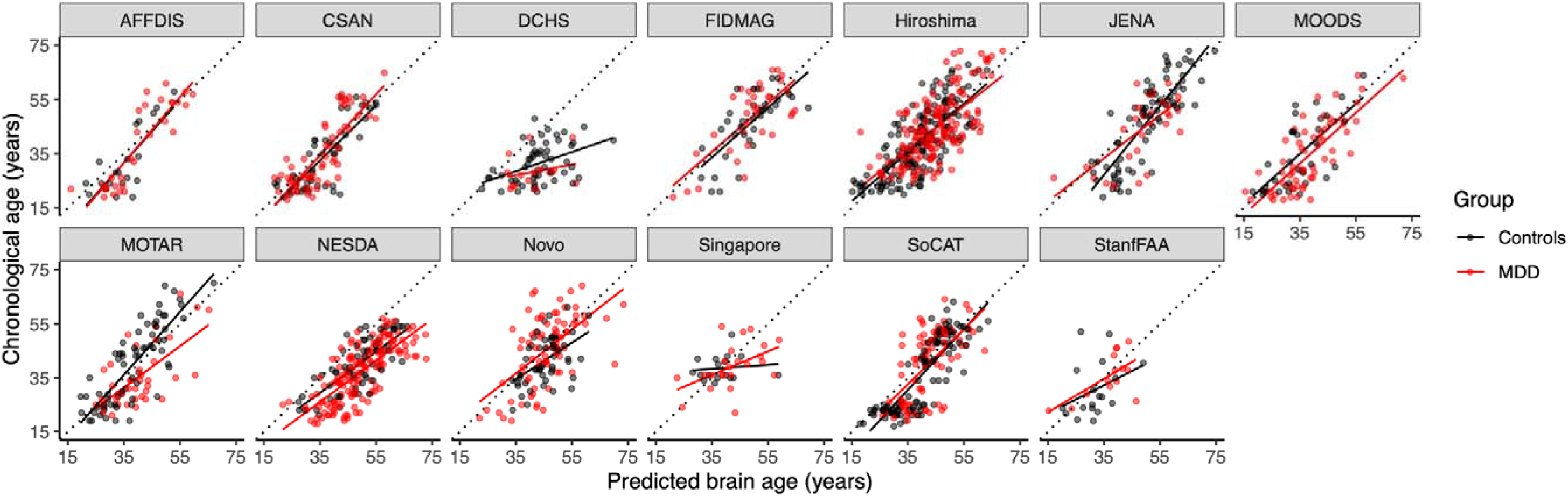
Brain age prediction using the ENIGMA algorithm in 13 new and unseen cohorts from 20 different scanners. Predicted brain age against chronological age per cohort. Separate regression lines are plotted for controls (black) and persons with depression (red). Diagonal dashed line reflects the line of identity (x=y).

### 3.3. Replication of higher brain age in depression

On average, depressed persons showed a significantly higher brain-PAD of +0.99 (SE 0.35) years (Cohen’s *d*□=□0.15, 95% CI: 0.04–0.25) compared with controls (*p*<0.01), **Figure 3**. No significant three-way interaction between diagnosis by age and by sex, nor significant two-way interactions (diagnosis by age or diagnosis by sex) were found. The supplementary meta-analytic approach resulted in a slightly higher but similar pooled brain-PAD of +1.20 years and associated effect size of Cohen’s *d*=0.19 between cases and controls (*I^2^*=52.6%, indicating moderate heterogeneity). Forest plots are depicted in **Supplementary Figure S2**.

**Figure 3.**
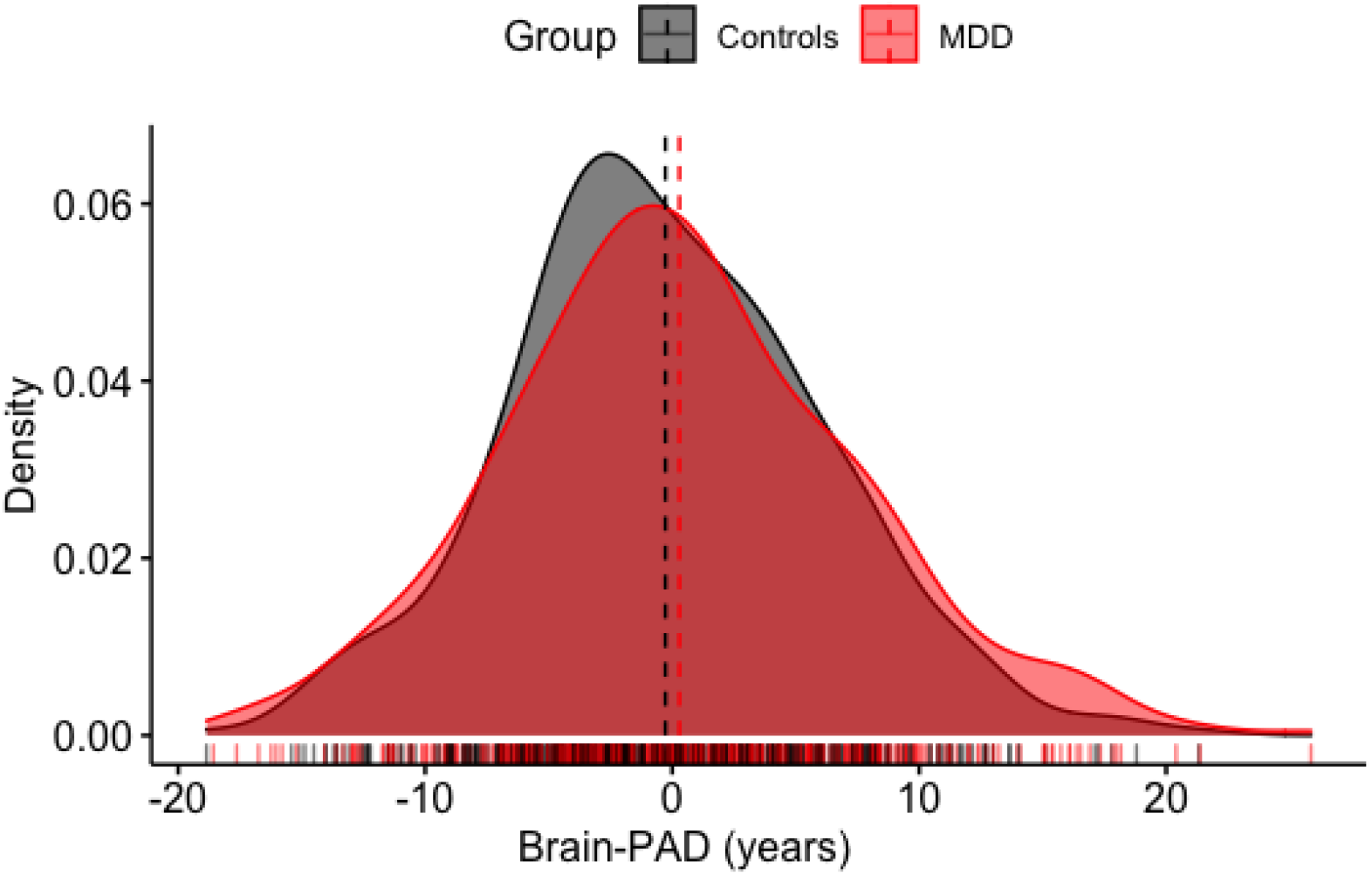
Replication of the brain age gap difference between controls and persons with depression. Brain-PAD (predicted brain age minus chronological age) in persons with major depressive disorder (MDD) and controls. Group-level analyses showed significantly higher brain-PAD in persons with MDD than controls in pooled samples of thirteen cohorts (*b*=0.99 years, *p*=0.003). The brain-PAD estimates are adjusted for chronological age, age^2^, sex and scanning site.

### 3.4. Patient group analyses and meta-regressions with moderators

No significant differences in brain-PAD were found between patient subgroups (recurrent versus first episode depression [b=0.00 years, p=0.99], AD-free versus AD-using patients [b=0.83 years, p=0.20], acute versus remitted depression [b=-1.07 years, p=0.43], or age of onset of depression in middle [b=0.55, p=0.49] or late adulthood [b=0.84, p=0.66] compared to early onset). The meta-regressions with sample size, mean age, proportion of females, proportion of first/recurrent episode patients, proportion of AD-free/AD-using patients, proportion of remitted/acute patients, field strength and performance accuracy metrics (MAE, R^2^) did not significantly moderate the Cohen’s d effect size estimates of brain-PAD (all QMp’s>0.05, **Supplementary Table S6**), but significant moderating effects of FreeSurfer version 5 (d=0.20, p=0.02) and version 6 (d=0.40, p=0.005) and Philips scanner vendor (d=0.50, p<0.0001) were found (**Figure S3** and **Figure S4** in the **Supplement**).

## 4. DISCUSSION

The current study replicated the finding that persons with depression reliably show older appearing brains, with a similar estimated gap and associated effect size (+1 year, Cohen’s d=0.14) as previously found in our largest mega-analysis of brain age in depression to date (+1.08 years, Cohen’s d=0.14) (Han et al., 2021a). While the generalization of our algorithm varied from cohort to cohort, pooled metrics were comparable to the performance accuracy found in the out-of-test samples in the original study. Importantly, post-hoc sensitivity analyses revealed that the exclusion of cohorts showing poor generalization did not change our replication findings (**Supplement**). In addition, a meta-analytic approach resulted in a highly similar pooled effect size (+1.20 years, Cohen’s d=0.19), providing robust evidence for significant but subtle age-related structural brain patterns in depression compared to controls.

The current multi-site replication study provides further evidence that the brain age gap in depression is an estimated +1 year (Cohen’s d=0.14), consistent with our previous mega-analysis in 19 other cohorts (Han et al., 2021a) and another meta-analysis including an additional 6 studies (Ballester et al., 2022). Taken together, the impact of depression on age-related structural brain differences thus seems to be rather subtle. However, it is important to note that the small, pooled effect size did not result from consistent small effects in each individual cohort, as can be seen from the forest plots of the meta-analyses in **Supplementary Figure S2**. Instead, the subtlety of the effect seemed to be driven by the fact that four of the cohorts showed larger positive effects (Cohen’s d’s ranging from 0.40 to 0.67, mean=0.50), whereas remaining cohorts showed no significant effects. However, **Supplementary Table S6** shows that effect sizes were not related to model generalization (i.e., small, or negative effect sizes were not only observed in cohorts showing poor performance accuracy). While negative *R^2^* observed in some cohorts can likely be explained by lower values of the cortical thickness features in those particular cohorts (**Supplementary Figure S1**), the inconsistency in effect sizes between cohorts may rather be due to other sources of variation unrelated to basic cohort or clinical characteristics such as first episode vs. recurrent, antidepressant free vs. antidepressant using or acute vs. remitted patients, as we did not observe any differences between these subgroups. The current study did however examine several potential technical sources of bias such as field strength, scanner vendor, and FreeSurfer version. In terms of scan technical moderators, we found that image acquisition with a Philips scanner (in contrast to Siemens or General Electric vendors) and FreeSurfer version for preprocessing images (v5 and v6, in contrast to v7) showed significant moderating effects on the effect size of the case-control difference in the brain age gap (**Supplementary Figures S3** and **S4**). While scanner manufacturer and FreeSurfer pipeline differences may potentially lead to (small) non-negligible differences in cortical thickness, surface area, and volume (Potvin et al., 2016), it is unlikely that effects would be differential in cases versus controls within the same cohorts. These effects were corrected for in the mega-analysis and it seems more plausible that other heterogeneous demographic, psychosocial, clinical, or biological cohort-specific characteristics, which we did not measure, coincided with the scanner vendor variable (i.e., biological sampling bias).

A recent systematic review, for example, suggests a role for epigenetic factors, and work investigating whether (genetic risk) for epigenetic aging contributes to the brain-PAD metric is underway in the ENIGMA consortium. While other literature suggests differential brain aging effects in older adults compared to middle-aged adults (i.e., only significantly higher brain-PAD in geriatric sample)(Christman et al., 2020), females and males (i.e., brain-PAD only associated with depressive severity in males)(Dunlop et al., 2021), or stage-dependent relationships with depression (i.e., only occurring at illness onset)(Han et al., 2021), we did not confirm this in the current study. Furthermore, detailed information on ethnicity, socioeconomic and psychosocial variance were not available and its impact on (the performance of the) brain age (prediction model) could not be evaluated in more detail here. However, an independent study including the NESDA cohort showed selectively older appearing brains in those with high *somatic* symptom severity (Han et al., 2021b). Future studies with more detailed (clinical) characterization (e.g., individual or clusters of depressive symptoms) are needed to gain more insight into which factors consistently contribute to the brain-PAD metric.

A major strength of this replication study is the harmonized approach of data preprocessing, quality checking, and brain age prediction algorithm across cohorts, potentially limiting the sources of bias that may stem from these decisions. This study is therefore a good example of the advantage of consortium efforts and collaborative team science. A note of caution is however due, since within individual cohorts, the case-control difference may not be consistent, present, or significant, also explaining the inconsistent findings across individual studies (Ballester et al., 2022). Unfortunately, due to a lack of harmonized clinical, demographic, and psychosocial information in consortia like ENIGMA MDD, we are limited in our ability to identify factors that could explain the variance in brain-PAD between cohorts. Finally, while the brain age predictions may be more accurate with higher-dimensional data from multimodal sources, it remains an open question whether models with improved performance accuracy show increased sensitivity in detecting subsequent associations with clinical psychopathology.

## 5. CONCLUSION

This replication study using data from 13 cohorts around the world confirmed our previous findings that persons with major depressive disorder show advanced brain aging compared to controls by approximately +1 year. Thus, two large-scale independent but harmonized mega-analyses across 32 cohorts and >3,400 patients and >2,800 controls show a reliable but subtle pattern of brain aging in adult depression. It is important to note that the small, pooled effect is not due to consistent small effects across cohorts but may be driven in part by the heterogeneity across scanning sites. Although we did not find a relation between basic patient properties and the effect size difference in the brain age gap, future work is needed to examine which clinical or biological characteristics may underlie the individual variation in the brain age gap.

## Data availability statement

The datasets for this manuscript are part of the ENIGMA MDD consortium and not publicly available. The used brain age prediction model is publicly available on www.photon-ai.com/enigma_brainage.

## Acknowledgements

We would like to thank all research participants, ENIGMA MDD working group contributors and collaborators for sharing their data and promoting team science.

## REFERENCES

Anderton, B.H., 2002. Ageing of the brain. Mech. Ageing Dev. 123, 811–817.

Baecker, L., Garcia-Dias, R., Vieira, S., Scarpazza, C., Mechelli, A., 2021. Machine learning for brain age prediction: Introduction to methods and clinical applications. EBioMedicine 72, 103600.

Ballester, P.L., Romano, M.T., de Azevedo Cardoso, T., Hassel, S., Strother, S.C., Kennedy, S.H., Frey, B.N., 2022. Brain age in mood and psychotic disorders: a systematic review and meta-analysis. Acta Psychiatr. Scand. 145, 42–55.

Christman, S., Bermudez, C., Hao, L., Landman, B.A., Boyd, B., Albert, K., Woodward, N., Shokouhi, S., Vega, J., Andrews, P., Taylor, W.D., 2020. Accelerated brain aging predicts impaired cognitive performance and greater disability in geriatric but not midlife adult depression. Transl. Psychiatry 10, 317.

Clausen, A.N., Fercho, K.A., Monsour, M., Disner, S., Salminen, L., Haswell, C.C., Rubright, E.C., Watts, A.A., Buckley, M.N., Maron-Katz, A., Sierk, A., Manthey, A., Suarez-Jimenez, B., Olatunji, B.O., Averill, C.L., Hofmann, D., Veltman, D.J., Olson, E.A., Li, G., Forster, G.L., Walter, H., Fitzgerald, J., Théberge, J., Simons, J.S., Bomyea, J.A., Frijling, J.L., Krystal, J.H., Baker, J.T., Phan, K.L., Ressler, K., Han, L.K.M., Nawijn, L., Lebois, L.A.M., Schmaal, L., Densmore, M., Shenton, M.E., van Zuiden, M., Stein, M., Fani, N., Simons, R.M., Neufeld, R.W.J., Lanius, R., van Rooij, S., Koch, S.B.J., Bonomo, S., Jovanovic, T., deRoon-Cassini, T., Ely, T.D., Magnotta, V.A., He, X., Abdallah, C.G., Etkin, A., Schmahl, C., Larson, C., Rosso, I.M., Blackford, J.U., Stevens, J.S., Daniels, J.K., Herzog, J., Kaufman, M.L., Olff, M., Davidson, R.J., Sponheim, S.R., Mueller, S.C., Straube, T., Zhu, X., Neria, Y., Baugh, L.A., Cole, J.H., Thompson, P.M., Morey, R.A., 2022. Assessment of brain age in posttraumatic stress disorder: Findings from the ENIGMA PTSD and brain age working groups. Brain Behav. 12, e2413.

Cole, J.H., Franke, K., 2017. Predicting Age Using Neuroimaging: Innovative Brain Ageing Biomarkers. Trends Neurosci. 40, 681–690.

Cole, J.H., Ritchie, S.J., Bastin, M.E., Valdés Hernández, M.C., Muñoz Maniega, S., Royle, N., Corley, J., Pattie, A., Harris, S.E., Zhang, Q., Wray, N.R., Redmond, P., Marioni, R.E., Starr, J.M., Cox, S.R., Wardlaw, J.M., Sharp, D.J., Deary, I.J., 2018. Brain age predicts mortality. Mol. Psychiatry 23, 1385–1392.

Constantinides, C., Han, L.K.M., Alloza, C., Antonucci, L., Arango, C., Ayesa-Arriola, R., Banaj, N., Bertolino, A., Borgwardt, S., Bruggemann, J., Bustillo, J., Bykhovski, O., Carr, V., Catts, S., Chung, Y.-C., Crespo-Facorro, B., Díaz-Caneja, C.M., Donohoe, G., Plessis, S.D., Edmond, J., Ehrlich, S., Emsley, R., Eyler, L.T., Fuentes-Claramonte, P., Georgiadis, F., Green, M., Guerrero-Pedraza, A., Ha, M., Hahn, T., Henskens, F.A., Holleran, L., Homan, S., Homan, P., Jahanshad, N., Janssen, J., Ji, E., Kaiser, S., Kaleda, V., Kim, M., Kim, W.-S., Kirschner, M., Kochunov, P., Kwak, Y.B., Kwon, J.S., Lebedeva, I., Liu, J., Mitchie, P., Michielse, S., Mothersill, D., Mowry, B., de la Foz, V.O.-G., Pantelis, C., Pergola, G., Piras, F., Pomarol-Clotet, E., Preda, A., Quidé, Y., Rasser, P.E., Rootes-Murdy, K., Salvador, R., Sangiuliano, M., Sarró, S., Schall, U., Schmidt, A., Scott, R.J., Selvaggi, P., Sim, K., Skoch, A., Spalletta, G., Spaniel, F., Thomopoulos, S.I., Tomecek, D., Tomyshev, A.S., Tordesillas-Gutiérrez, D., van Amelsvoort, T., Vázquez-Bourgon, J., Vecchio, D., Voineskos, A., Weickert, C.S., Weickert, T., Thompson, P.M., Schmaal, L., van Erp, T.G.M., Turner, J., Cole, J.H., Dima, D., Walton, E., 2022. Brain ageing in schizophrenia: evidence from 26 international cohorts via the ENIGMA Schizophrenia consortium. bioRxiv. https://doi.org/10.1101/2022.01.10.21267840

Dunlop, K., Victoria, L.W., Downar, J., Gunning, F.M., Liston, C., 2021. Accelerated brain aging predicts impulsivity and symptom severity in depression. Neuropsychopharmacology 46, 911–919.

Han, L.K.M., Dinga, R., Hahn, T., Ching, C.R.K., Eyler, L.T., Aftanas, L., Aghajani, M., Aleman, A., Baune, B.T., Berger, K., Brak, I., Filho, G.B., Carballedo, A., Connolly, C.G., Couvy-Duchesne, B., Cullen, K.R., Dannlowski, U., Davey, C.G., Dima, D., Duran, F.L.S., Enneking, V., Filimonova, E., Frenzel, S., Frodl, T., Fu, C.H.Y., Godlewska, B.R., Gotlib, I.H., Grabe, H.J., Groenewold, N.A., Grotegerd, D., Gruber, O., Hall, G.B., Harrison, B.J., Hatton, S.N., Hermesdorf, M., Hickie, I.B., Ho, T.C., Hosten, N., Jansen, A., Kähler, C., Kircher, T., Klimes-Dougan, B., Krämer, B., Krug, A., Lagopoulos, J., Leenings, R., MacMaster, F.P., MacQueen, G., McIntosh, A., McLellan, Q., McMahon, K.L., Medland, S.E., Mueller, B.A., Mwangi, B., Osipov, E., Portella, M.J., Pozzi, E., Reneman, L., Repple, J., Rosa, P.G.P., Sacchet, M.D., Sämann, P.G., Schnell, K., Schrantee, A., Simulionyte, E., Soares, J.C., Sommer, J., Stein, D.J., Steinsträter, O., Strike, L.T., Thomopoulos, S.I., van Tol, M.-J., Veer, I.M., Vermeiren, R.R.J.M., Walter, H., van der Wee, N.J.A., van der Werff, S.J.A., Whalley, H., Winter, N.R., Wittfeld, K., Wright, M.J., Wu, M.-J., Völzke, H., Yang, T.T., Zannias, V., de Zubicaray, G.I., Zunta-Soares, G.B., Abé, C., Alda, M., Andreassen, O.A., Bøen, E., Bonnin, C.M., Canales-Rodriguez, E.J., Cannon, D., Caseras, X., Chaim-Avancini, T.M., Elvsåshagen, T., Favre, P., Foley, S.F., Fullerton, J.M., Goikolea, J.M., Haarman, B.C.M., Hajek, T., Henry, C., Houenou, J., Howells, F.M., Ingvar, M., Kuplicki, R., Lafer, B., Landén, M., Machado-Vieira, R., Malt, U.F., McDonald, C., Mitchell, P.B., Nabulsi, L., Otaduy, M.C.G., Overs, B.J., Polosan, M., Pomarol-Clotet, E., Radua, J., Rive, M.M., Roberts, G., Ruhe, H.G., Salvador, R., Sarró, S., Satterthwaite, T.D., Savitz, J., Schene, A.H., Schofield, P.R., Serpa, M.H., Sim, K., Soeiro-de-Souza, M.G., Sutherland, A.N., Temmingh, H.S., Timmons, G.M., Uhlmann, A., Vieta, E., Wolf, D.H., Zanetti, M.V., Jahanshad, N., Thompson, P.M., Veltman, D.J., Penninx, B.W.J.H., Marquand, A.F., Cole, J.H., Schmaal, L., 2021a. Brain aging in major depressive disorder: results from the ENIGMA major depressive disorder working group. Mol. Psychiatry 26, 5124–5139.

Han, L.K.M., Schnack, H.G., Brouwer, R.M., Veltman, D.J., van der Wee, N.J.A., van Tol, M.-J., Aghajani, M., Penninx, B.W.J.H., 2021b. Contributing factors to advanced brain aging in depression and anxiety disorders. Transl. Psychiatry 11, 402.

Han, S., Chen, Y., Zheng, R., Li, S., Jiang, Y., Wang, C., Fang, K., Yang, Z., Liu, L., Zhou, B., Wei, Y., Pang, J., Li, H., Zhang, Y., Cheng, J., 2021. The stage-specifically accelerated brain aging in never-treated first-episode patients with depression. Hum. Brain Mapp. 42, 3656–3666.

Lombardi, A., Monaco, A., Donvito, G., Amoroso, N., Bellotti, R., Tangaro, S., 2020. Brain Age Prediction With Morphological Features Using Deep Neural Networks: Results From Predictive Analytic Competition 2019. Front. Psychiatry 11, 619629.

Penninx, B., 2017. Depression and its Somatic Consequences: Allostatic Load as the Connecting Link. Eur. Psychiatry 41, S62–S62.

Potvin, O., Mouiha, A., Dieumegarde, L., Duchesne, S., 2016. P4-163: Effects of scanner manufacturer and strength on cortical surfaces, thicknesses and volumes in the aging brain. Alzheimers. Dement. 12, P1077–P1079.

Wrigglesworth, J., Ward, P., Harding, I.H., Nilaweera, D., Wu, Z., Woods, R.L., Ryan, J., 2021. Factors associated with brain ageing - a systematic review. BMC Neurol. 21, 312.

