## supplement for "A Large-Scale ENIGMA Multisite Replication Study of Brain Age in Depression"

**SUPPLEMENTARY MATERIALS**

**Supplementary Tables**

- **Supplementary Table S1.** ENIGMA - Major Depressive Disorder Working Group Sample Characteristics by cohort.
- **Supplementary Table S2.** ENIGMA - Major Depressive Disorder Working Group Image acquisition and processing by cohort.
- **Supplementary Table S3.** ENIGMA - Major Depressive Disorder Working Group Instrument for diagnosing Major Depressive Disorder and exclusion criteria by cohort.
- **Supplementary Table S4.** Generalizability to independent patient cohorts.
- **Supplementary Table S5.** Sensitivity analysis: pooled performance accuracy and case-control difference in brain-PAD.
- **Supplementary Table S6.** Results from the meta-regressions with moderators.

**Supplementary Figures**

- **Supplementary Figure S1.** Mean cortical thickness values per cohort.
- **Supplementary Figure S2.** Forest plots of case-control comparisons of brain-PAD. Panel **(A)** displays Cohen's d effect size estimates and panel **(B)** shows the estimated brain-PAD difference in years within each individual cohort.
- **Supplementary Figure S3.** Forest plot with three subgroups of FreeSurfer version used for preprocessing.
- **Supplementary Figure S4.** Forest plot with three subgroups of scanner vendors used for image acquisition.

**Supplementary Table S1.** Participants characteristics per cohort.

| **Characteristic** | **AFFDIS**, N = 49*^1^* | **CSAN**, N = 107*^1^* | **DCHS**, N = 70*^1^* | **FIDMAG**, N = 69*^1^* | **Hiroshima**, N = 315*^1^* | **TiPs**, N = 105*^1^* | **MOODS,** N = 96^1^ | **MOTAR**, N = 108*^1^* | **NESDA**, N = 219*^1^* | **Novo**, N = 128*^1^* | **Singapore**, N = 40*^1^* | **SoCAT**, N = 179*^1^* | **StanfFAA**, N = 32*^1^* |
| --- | --- | --- | --- | --- | --- | --- | --- | --- | --- | --- | --- | --- | --- |
| Controls | 20 (41%) | 49 (46%) | 54 (77%) | 34 (49%) | 167 (53%) | 75 (71%) | 32 (33%) | 68 (63%) | 65 (30%) | 52 (41%) | 17 (42%) | 100 (56%) | 18 (56%) |
| MDD | 29 (59%) | 58 (54%) | 16 (23%) | 35 (51%) | 148 (47%) | 30 (29%) | 64 (67%) | 40 (37%) | 154 (70%) | 76 (59%) | 23 (57%) | 79 (44%) | 14 (44%) |
| Chronological age (years) | 35.73 ± 14.35 (19.00-61.00) | 34.25 ± 12.61 (18.00-65.00) | 30.69 ± 7.05 (21.00-48.00) | 47.12 ± 12.30 (19.00-66.00) | 41.82 ± 12.30 (20.00-73.00) | 46.15 ± 14.55 (19.00-73.00) | 34.54 ± 12.54 (18.00-64.00) | 38.34 ± 13.55 (19.00-70.00) | 38.11 ± 10.35 (18.00-57.00) | 43.92 ± 11.57 (19.00-69.00) | 38.98 ± 6.94 (22.00-54.00) | 37.85 ± 13.37 (17.00-64.00) | 32.71 ± 9.71 (18.93-52.07) |
| Predicted brain age (years) | 38.34 ± 10.22 (16.05-59.50) | 35.60 ± 9.68 (14.78-57.91) | 42.78 ± 8.74 (22.78-69.78) | 48.66 ± 9.82 (21.11-69.37) | 41.90 ± 10.31 (15.16-68.82) | 49.98 ± 10.17 (17.13-74.60) | 37.11 ± 10.01 (15.51-71.90) | 39.02 ± 10.02 (19.48-67.00) | 49.52 ± 10.17 (21.77-75.66) | 45.75 ± 8.19 (22.04-73.39) | 40.70 ± 8.60 (22.55-58.94) | 40.98 ± 9.26 (17.91-63.62) | 33.78 ± 7.93 (15.22-49.07) |
| Brain-PAD (years) | 2.60 ± 7.68 (-14.90-19.05) | 1.35 ± 7.44 (-14.41-22.20) | 12.09 ± 8.63 (-6.36-32.40) | 1.54 ± 8.46 (-15.02-20.57) | 0.08 ± 8.61 (-24.94-21.57) | 3.83 ± 8.92 (-14.20-24.09) | 2.57 ± 8.62 (-19.57-20.79) | 0.67 ± 9.12 (-19.46-24.34) | 11.41 ± 7.40 (-6.36-30.89) | 1.83 ± 9.78 (-25.19-30.29) | 1.72 ± 8.64 (-15.52-22.71) | 3.13 ± 8.72 (-18.01-24.14) | 1.07 ± 8.86 (-25.31-20.24) |
| Sex Male | 26 (53%) | 34 (32%) | - | 26 (38%) | 140 (44%) | 53 (50%) | 31 (32%) | 53 (49%) | 74 (34%) | 44 (34%) | 21 (52%) | 18 (10%) | - |
| Female | 23 (47%) | 73 (68%) | 70 (100%) | 43 (62%) | 175 (56%) | 52 (50%) | 65 (68%) | 55 (51%) | 145 (66%) | 84 (66%) | 19 (48%) | 161 (90%) | 32 (100%) |
| Recurrent status Control | 20 (41%) | 49 (46%) | NA | 34 (51%) | 167 (53%) | 75 (71%) | 32 (33%) | 68 (63%) | 65 (30%) | 52 (41%) | 17 (42%) | 100 (56%) | 18 (56%) |
| First | 2 (4.1%) | 14 (13%) | NA | 11 (16%) | 73 (23%) | 7 (6.7%) | 32 (33%) | 29 (27%) | 67 (31%) | 36 (28%) | 8 (20%) | 19 (11%) | - |
| Recurrent | 27 (55%) | 44 (41%) | NA | 22 (33%) | 73 (23%) | 23 (22%) | 32 (33%) | 11 (10%) | 87 (40%) | 40 (31%) | 15 (38%) | 60 (34%) | 14 (44%) |
| AD status Control | 20 (41%) | 49 (46%) | NA | 34 (50%) | 167 (53%) | 74 (70%) | 32 (33%) | 68 (63%) | 65 (30%) | 52 (41%) | 17 (42%) | 100 (56%) | 18 (56%) |
| AD-free | 1 (2.0%) | 30 (28%) | NA | 4 (5.9%) | 10 (3.2%) | 12 (11%) | 64% (67%) | 40 (37%) | 98 (45%) | 51 (40%) | 5 (12%) | 41 (23%) | 11 (34%) |
| AD-using | 28 (57%) | 28 (26%) | NA | 30 (44%) | 136 (43%) | 19 (18%) | 0 (0%) | - | 56 (26%) | 25 (20%) | 18 (45%) | 38 (21%) | 3 (9.4%) |
| Remitted status Control | 20 (41%) | 49 (46%) | NA | 34 (49%) | 167 (53%) | NA | 32 (33%) | 68 (63%) | 65 (30%) | 52 (41%) | 17 (100%) | 100 (56%) | 18 (56%) |
| Remitted | - | - | NA | 1 (1.4%) | - | NA | 0 (0%) | - | - | 7 (5.5%) | - | 33 (18%) | - |
| Acute | 29 (59%) | 58 (54%) | NA | 34 (49%) | 148 (47%) | NA | 64 (67%) | 40 (37%) | 154 (70%) | 69 (54%) | - | 46 (26%) | 14 (44%) |
| FreeSurfer version | 5 | 7 | 5 | 6 | 5 | 5 | 6 | 6 | 5 | 5 | 5 | 7 | 5 |
| Field strength | 3T | 3T | 3T | 1.5T | 3T | 3T | 3T | 3T | 3T | 3T | 3T | 3T | 3T |
| Scanner vendor | Siemens | Siemens | Siemens | GE | GE/Siemens | Siemens | Philips | Philips | Philips | GE | Philips | Siemens | GE |
| *^1^* n (%); Mean ± SD (Minimum-Maximum) | | | | | | | | | | | | | |

**Supplementary Table S2.** ENIGMA - Major Depressive Disorder Working Group Image acquisition and processing by cohort.

| **Cohort** | **Country** | **Scanner type** | **Sequence T1** | **FreeSurfer version** | **Slice orientation** | **Operating system** |
| --- | --- | --- | --- | --- | --- | --- |
| **AFFDIS** | Germany | 3T Siemens Magnetom TrioTim | 3D T1 (176 slices; TR = 2250 ms; TE = 3.26 ms; FOV 256; voxel size 1X1X1mm) | 5.3 | Sagittal | Linux CentOS |
| **CSAN** | Sweden | 3T Siemens MAGNETOM PRISMA | 3D T1: MP-RAGE (TR = 2300 ms, TE = 2.34 ms, FOV 250 × 250 mm, voxel size = 0.9 × 0.868 × 0.868 mm, flip angle = 8°) | 7.2 | Sagittal | Ubuntu |
| **DCHS** | South Africa | 3T Siemens Skyra | 3D T1: multi-echo MPRAGE, voxel size 1 mm x 1mm x 1.5mm, TR = 2530 ms, TE = 1.69 x 3.55 x 5.41 x 7.27ms, FOV: 256x256mm, flip angle = 7° | 5.3 | Sagittal | Linux-centos6_x86_64 |
| **FIDMAG** | Spain | 1.5T GE Signa | 3D T1: matrix size = 512 × 512, 180 contiguous axial slices, voxel resolution = 0.47 × 0.47 × 1mm, no slice gap, TE = 3.93ms, TR = 2000ms and inversion time (TI) = 710ms, flip angle = 15 degrees | 6.0 | Axial | Linux-centos6_x86_64 |
| **Hiroshima1** | Japan | 3T GE Signa HDxt | 3D T1: 256x256x256 matrix of 1x1x1mm voxels (GE: SPGR, 184 slices) | 5.3 | Sagittal | Linux_Ubuntu_18.04 |
| **Hiroshima2** | Japan | 3T SIEMENS MAGNETOM Spectra | 3D T1: 256x256x256 matrix of 1x1x1mm voxels (Siemens: ADNI MPRAGE (tfl), GRAPPA, 192 slices) | 5.3 | Sagittal | Linux_Ubuntu_18.04 |
| **MOODS/DEP-ARREST CLIN** | France | 3T Philips Achieva | 3D T1: TR=7, TE=3.5, FOV=352x352x180, Flip angle=8 degrees, number of slices : 180 slices, Slice gap 1 mm, voxel size: 0.8x0.8x1 | 6.0 | Transverse (Axial) | CentOS Linux 7 |
| **MOTAR** | Netherlands | 3T Philips Achieva | 3D T1: 32-channel head coil at the Spinoza centre. 3D Turbo Field Echo (TFE), TR = 8.1 ms, TE = 3.7 ms, flip angle = 8 degrees, matrix size = 240 x 240, 1 mm^3^ isotropic voxels. | 6.0 | Transverse (Axial) | SHARK HPC, Linux environment |
| **NESDA** | Netherlands | 3T Phillips Achieva/Intera | 3D T1: gradient-echo TR=9 msec; TE=3.5 msec; flip angle 8º, FOV = 256 mm; matrix: 25x62x56; in plane voxel size = 1 mm × 1 mm x 1 mm; 170 slices. | 5.0 | Sagittal | SHARK HPC, Linux environment |
| **Novosibirsk** | Russia | 3T GE Discovery™ MR750w | 3D T1: fast spin gradient echo sequence (FSPGR BRAVO), repetition time = 9.5 ms, echo time = 3.7 ms, flip angle = 3°, acquired over a field of view of 256 (feet-head [FH]) × 256 (anterior-posterior [AP]) × 188 (rightleft [RL]) mm, reconstructed to cubic voxels of 1 mm × 1 mm × 1 mm | 5.3 | Sagittal | OS X 10.10 |
| **Singapore** | Singapore | 3T Philips Achieva | 3D T1: MP-RAGE (magnetisation-prepared rapid acquisition with a gradient echo) volumetric scans (TR/TE/TI/flip angle 8.4/3.8/3000/8; matrix 256x204; FOV 240mm2) with axial orientation (reformatted to coronal) | 5.3 | Axial | Linux_Ubuntu12.04_6 4 |
| **SoCAT** | Turkey | 3T Siemens Verio,Numaris/4,Syngo MR B17 | 3D T1: MP-Rage/axial plane; TR=1900 msec; TE=3.4 msec; Flip angle=15°; Voxel size 1 mm x 1 mm x 1 mm | 7.1 | Axial | freesurfer-linux-centos7_x86_64-7.1.1 |
| **StanfFAA** | United States of America | 3T GE Discovery MR750 | 3D T1: spoiled gradient echo (SPGR) pulse sequence (186 sagittal slices; resolution = 0.9 mm isotropic; flip angle = 12°; repetition time [TR] = 6,240 ms; echo time [TE] = 2.34 ms) | 5.3 | Sagittal | Centos6_x86_64, Linux-based HPC |
| **TiPs** | Germany | 3T Siemens MAGNETOM Prisma_fit | 3D T1: MP-RAGE sequence: TR 2300 ms, TE 3.03 ms, α 9°, 192 contiguous sagittal slices, in-plane field of view 256 mm, voxel resolution 1Å~1Å~1 mm; acquisition time 5:21 min | 5.3 | Sagittal | Linux |

**Supplementary Table S3.** ENIGMA - Major Depressive Disorder Working Group Instrument for diagnosing Major Depressive Disorder and exclusion criteria by cohort.

| **Cohort** | **Diagnostic instrument** | **Sample characteristics/Inclusion criteria** | **Exclusion criteria** |
| --- | --- | --- | --- |
| **AFFDIS** | ICD-10, DSM-IV criteria | MDD subjects were currently depressed and in day program or inpatient | All subjects exclusion criteria: current or history of neurological disorder or brain injury, current substance abuse or dependence (not including nicotine), pregnancy, MRI contraindications, inability to give consent. MDD specific: comorbid psychiatric diagnosis. Healthy control specific: current or history of psychiatric diagnosis. |
| **CSAN** | MINI | MDD subjects were currently depressed meeting MINI criteria for depression; comorbid anxiety disorders are allowed; mood-congruent psychotic symptoms allowed. | Current MDD: a current DSM-5 diagnosis of substance use disorder, except nicotine; a psychotic disorder, except depression with mood-congruent psychotic features; new antidepressant medication during the month before study participation (two months for fluoxetine); change of the dose of psychotropic medications over the last month (antidepressant and antipsychotic medication) or the last two months (mood stabilizers and anticonvulsants). |
| **DCHS** | MINI | DCHS is a South African birth cohort study including women over the age of 18 years, who were between 20 and 28 weeks pregnant, presented at either of two recruitment clinics, and were able to give written informed consent. MRI was obtained in a subsample of mothers (at least 18 months after birth) with a lifetime MINI diagnosis of MDD and in controls without MINI diagnosis. | In the DCHS MRI substudy mothers were excluded for: loss of consciousness longer than 30 minutes; inability to speak English; current/lifetime alcohol and/or substance dependence or abuse; psychopathology other than PTSD and/or MDD; traumatic brain injury; standard MRI exclusion criteria. |
| **FIDMAG** | DSM-IV-TR criteria | MDD subjects were currently depressed (HDRS >= 17, only 1 patient was in remission), right-handed, age 18-65 | Patients were excluded (i) if they were left-handed; (ii) if they were younger than 18 or older than 65 years; (iii) if they had a history of brain trauma or neurological disease; (iv) if they had shown alcohol/ substance abuse within 12 months prior to participation; and (v) if they had undergone electroconvulsive therapy in the previous 12 months. |
| **Hiroshima1** | MINI | MDD subjects were recruited from local clinics, 20-80 years. Controls were recruited from local community by advertising in local papers. | MDD patients: comorbid psychiatric disorders other than MDD, Control subjects: any history of psychiatric disorder |
| **Hiroshima2** | MINI | MDD subjects were recruited from local clinics, 20-80 years. Controls were recruited from local community by advertising in local papers. | MDD patients: comorbid psychiatric disorders other than MDD, Control subjects: any history of psychiatric disorder |
| **MOODS / DEP-ARREST CLIN** | MINI, DSM-IV criteria | MDD subjects were currently depressed meeting MINI criteria for depression (HDRS>=17); Free of antidepressant use at least one month before the study. Control subjects were included based on the absence of current or past mental disorders or somatic conditions, particularly nasal polyposis and chronic or acute sinusitis or rhinitis; age 18-65. | Patients suffering from bipolar disorder, psychotic disorder, eating disorder, and addictions, according to the DSM-5 criteria, or from nasal polyposis, chronic or acute sinusitis, chronic or acute rhinitis or pregnancy or breastfeeding, were not included. HCs were included based on the absence of current or past mental disorders or somatic conditions, particularly nasal polyposis and chronic or acute sinusitis or rhinitis |
| **MOTAR** | CIDI, DSM-IV criteria | Inclusion criteria of the patient sample include: having a current depressive disorder (major depressive disorder) or anxiety disorder (social phobia, generalized anxiety disorder, panic disorder with or without agoraphobia) as ascertained by the Diagnostic and Statistical Manual of Mental Disorders – Fourth edition (DSM-IV) algorithms with the CIDI (Composite International Diagnostic Interview) and being aged between 18 and 70 years. | Severe internal or neurological disorders, MRI contraindications, use of antidepressants or other psychoactive medication (with the exception of stable benzodiazepine use for patients), lifetime diagnosis of psychotic disorders, bipolar disorder or personality disporders and dependence on drugs or alcohol. |
| **NESDA** | CIDI, DSM-IV criteria | DSM-4 based diagnosis of MDD (6 month recency), using CIDI interview. 93 (60%) MDD patients have a comorbid ANX diagnosis. Age range 18-65 | |
| **Novo** | MINI, SCID, ICD-10 | MDD Patients: 100% hospital-based, outpatients: 0%, general population: 0% | MDD subjects: Presence of axis-I disorders other than MDD, panic disorder, social anxiety disorder, or generalized anxiety disorder and any use of psychotropic medication other than stable use of SSRIs or infrequent benzodiazepine use; age 18 or below; alcohol or substance abuse/dependence within 6 months of study participation; current major medical problems. Control subjects: age over 65; any current or former psychiatric disorder. Both groups: MRI contra-indications. |
| **Singapore** | SCID | Inclusion: 1) DSM IV dx of MDD (Patients) 2) Age: 21-65 3) English speaking 4) Provision of informed written consent | Exclusion criteria 1) History of significant head injury 2)Neurological diseases such as epilepsy, cerebrovascular accident 3) Impaired thyroid function 4) Steroid use 5) DSM IV alcohol or substance use or dependence 6) Contraindications to MRI (e.g. pacemaker, orbital foreign body, recent surgery/procedure with metallic devices/implants deployed) using standard MRI Request Form from NNI 7)Pregnant women 8) Claustrophobia |
| **SoCAT** | SCID | Inclusion criteria: DSM IV dx for mdd patients Age: 18-65 right-handed currently depressed or remitted; Control subjects: any history of psychiatric disorder | Exclusion criteria 1) History of significant head injury 2)Neurological diseases such as epilepsy, cerebrovascular accident 3)Other diagnoses on Axis I disorders4) |
| **StanfFAA** | SCID | Community-based DSM-diagnosed sample | MDD subjects: presence of axis-I disorders other than MDD, anxiety and eating disorders . Control subjects: control individuals did not meet diagnostic criteria for any current psychiatric. Both groups: alcohol / substance abuse or dependence within six months prior to MRI scanning, history of head trauma with loss of consciousness > 5 min, aneurysm, or any neurological or metabolic disorders that require ongoing medication or that may affect the central nervous system (including thyroid disease, diabetes, epilepsy or other seizures, or multiple sclerosis), MRI contraindications, or bad MRI data (e.g., extreme movement). |
| **TiPs** | SCID | Psychiatric inpatients and tinnitus patients with MDD or a disorder of the depressive spectrum (also adjustment disorders as pointed out in the data table); psychiatrically healthy controls were derived from community and tinnitus patients | MDD subjects: presence of axis-I disorders other than MDD or adjustment disorders. Control subjects: no Axis-I diagnosis, no medication use. Exclusion criteria for all subjects included history of neurological disease (e.g. tumour, head trauma, epilepsy) or untreated internal medical condtitions, intellectual and/or developmental disability. Only German native speakers were allowed to participate. |

**Supplementary Table S4.** Generalizability to independent patient cohorts.

|  | ***r*** | ***R^2^*** | **MAE** | **SD** | **_w_MAE** | **RMSE** | **Brain-PAD** | **SD** |
| --- | --- | --- | --- | --- | --- | --- | --- | --- |
| **Pooled** | 0.66 | 0.29 | 6.65 | 6.26 | 0.15 | 10.68 | 4.12 | 9.86 |
| **AFFDIS** | 0.88 | 0.72 | 6.81 | 4.12 | 0.16 | 7.92 | 2.11 | 7.77 |
| **CSAN** | 0.82 | 0.67 | 6.18 | 4.15 | 0.13 | 7.42 | 0.55 | 7.47 |
| **DCHS** | 0.25 | -6.25 | 13.63 | 7.79 | 0.68 | 15.58 | 13.19 | 8.57 |
| **FIDMAG** | 0.73 | 0.53 | 7.26 | 5.00 | 0.15 | 8.78 | 0.69 | 8.88 |
| **Hiroshima** | 0.67 | 0.42 | 7.31 | 5.36 | 0.15 | 9.05 | 0.32 | 9.08 |
| **TiPs** | 0.68 | 0.29 | 7.91 | 6.28 | 0.2 | 10.03 | 4.45 | 9.14 |
| **MOODS** | 0.78 | 0.51 | 7.14 | 4.95 | 0.15 | 8.67 | 3.82 | 7.84 |
| **MOTAR** | 0.66 | 0.22 | 8.02 | 5.82 | 0.17 | 9.87 | 4.79 | 8.74 |
| **NESDA** | 0.75 | -0.92 | 12.7 | 6.96 | 0.32 | 14.47 | 12.43 | 7.44 |
| **Novo** | 0.58 | 0.32 | 8.32 | 6.69 | 0.16 | 10.64 | 0.41 | 10.71 |
| **Singapore** | 0.49 | -0.30 | 7.23 | 5.99 | 0.22 | 9.31 | 3.12 | 8.97 |
| **SoCAT** | 0.62 | 0.36 | 8.55 | 5.73 | 0.19 | 10.27 | 2.25 | 10.09 |
| **StanfFAA** | 0.63 | 0.24 | 5.29 | 4.93 | 0.20 | 7.11 | 0.98 | 7.31 |

*r*, Pearson’s correlation coefficient; *R^2^*, explained variance; MAE, mean absolute error; ; SD, standard deviation; _w_MAE, weighted MAE ($MAE\div age range of sample$); RMSE, root mean squared error; brain-PAD, brain-predicted age difference (predicted brain age minus chronological age).

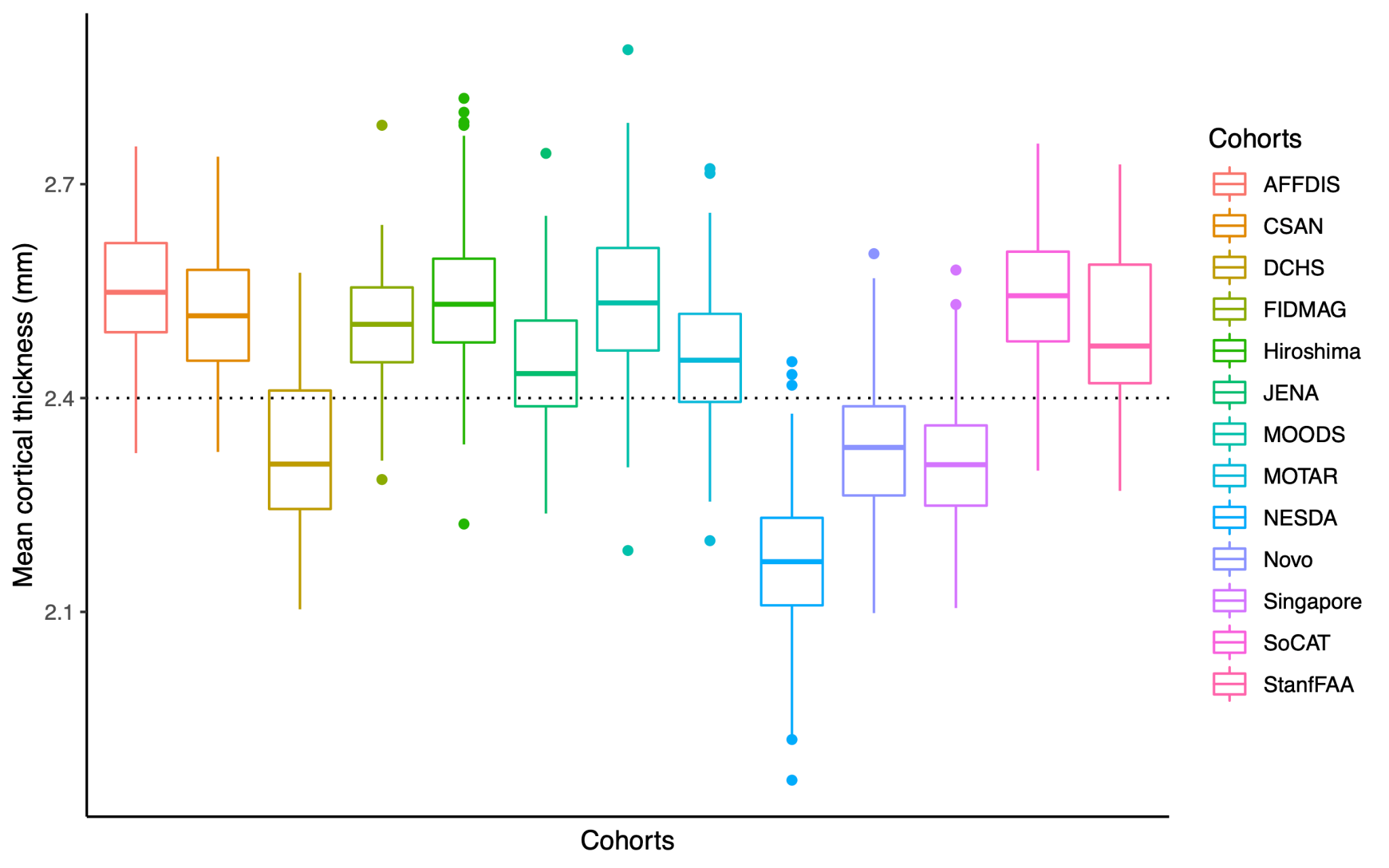

**Supplementary Figure S1.** Mean cortical thickness values per cohort. The dotted line indicates the grand mean (mean cortical thickness of 2.44 mm) of the sample. Cohorts with a negative deviation from the grand mean (i.e., DCHS [-0.11 mm], NESDA [-0.27 mm], Novo [-0.11 mm], Singapore [-0.13 mm]) also showed a negative *R^2^* between chronological age and predicted brain age (*R^2^’s* of -3.14, -0.35, -0.02, and -2.02, respectively).

*Sensitivity analyses*

Given that the generalizability of the brain age prediction model varied from cohort to cohort, we also performed sensitivity analyses by only including: **(1)** cohorts for which the correlation coefficient *r* between predicted brain age and chronological age was >.48, and **(2)** cohorts for which the explained variance *R^2^* was >.47. These arbitrary cut-offs were chosen to reflect cohorts showing **(1)** moderate-to-high correlation, but a regression intercept shift was observed and **(2)** both moderate-to-high correlation and explained variance, with limited shift of the regression intercept. The pooled performance accuracy for these subgroups are compared in **Supplementary Table S5**. In addition, **Supplementary Table S5** displays the Cohen’s *d* effect sizes for **(1)** the case-control comparison of brain-PAD in all thirteen cohorts (N=751 controls and N=766 persons with MDD); **(2)** the cohorts showing moderate-to-high correlation resulting in N=662 controls and N=713 persons with MDD from ten cohorts; and **(3)** cohorts showing moderate-to-high explained variance resulting in N=513 controls and N=419 persons with MDD from eight cohorts. All case-control comparisons showed similar estimated brain-PAD differences (b’s [in years] 0.99, 0.93, 1.01, respectively) and significantly higher brain-PAD in depression (Cohen’s *d*’s 0.15 vs. 0.14 vs. 0.16, respectively).

**Supplementary Table S5.** Sensitivity analysis: pooled performance accuracy and case-control difference in brain-PAD.

|  | **# Cohorts** | **r** | **R^2^** | **MAE** | **SD** | **_w_MAE** | **Brain-PAD** | **SD** | **b  (in years)** | **p** | **Cohen's d** | **se** | **95% CI lower** | **95% CI upper** |
| --- | --- | --- | --- | --- | --- | --- | --- | --- | --- | --- | --- | --- | --- | --- |
| **All controls** | **13** | 0.73 | 0.47 | 7.51 | 5.59 | 0.13 | 2.95 | 8.89 | 0.99 | 0.003 | 0.15 | 0.05 | 0.04 | 0.25 |
| **Moderate-to-high correlation** | **10** | 0.77 | 0.55 | 7.15 | 5.17 | 0.13 | 2.36 | 8.51 | 0.93 | 0.005 | 0.14 | 0.05 | 0.03 | 0.24 |
| **Moderate-to-high explained variance** | **8** | 0.80 | 0.62 | 6.89 | 4.98 | 0.12 | 1.42 | 8.39 | 1.01 | 0.006 | 0.16 | 0.06 | 0.04 | 0.28 |

Performance metrics were calculated in controls. Moderate-to-high correlation is defined as *r>*0.48 between predicted brain age and chronological age. Moderate-to-high explained variance is defined as *R^2^*>0.49. Regression coefficient b is obtained through a linear mixed model with fixed effects for age, age^2^, and sex and random effects for scanning site.

**META-ANALYTIC APPROACH**

*Methods*

A meta-analytical framework was used to compare the individual effect size estimates and their level of precision in each cohort. Meta-analysis models were fitted using the restricted maximum likelihood method (REML). In addition to meta-analyzed Cohen’s *d* effect size estimates and *b* brain age gap differences, heterogeneity scores (I^2^) were also calculated. This score indicates the total variance in effect size that can be explained by heterogeneity alone. Thus, lower values of I^2^ indicate lower variance in the effect size estimation across studies. Separate meta-regression analyses were used to test whether cohort characteristics explained a significant proportion of the variance in effect sizes between cohorts in the meta-analysis. Specifically, sample size, mean age, proportion of females, proportion of first episode/recurrent episode patients, proportion of antidepressant free/antidepressant using patients, proportion of remitted/acute patients, field strength of MR images, FreeSurfer version used for image processing, scanner vendor, and performance accuracy metrics (R^2^ and MAE) were included as a fixed effect predictor in meta-regression models.

*Results*

The meta-analytic approach resulted in a slightly higher but similar pooled brain age gap of +1.25 years and associated effect size of Cohen’s *d*=0.19 between cases and controls (*I^2^*=52.6%, indicating moderate heterogeneity), forest plots are depicted in **Figure S2**. The meta-regressions with sample size, mean age, proportion of females, proportion of first episode/recurrent episode patients, proportion of antidepressant free/antidepressant using patients, proportion of remitted/acute patients, field strength and performance accuracy metrics (MAE, R^2^) did not significantly moderate Cohen’s d effect size estimates of brain-PAD (all QMp’s>0.05), but significant moderating effects of FreeSurfer version 5 (d=0.23, p=0.02) and version 6 (d=0.41, p=0.009) and Philips scanner vendor (d=0.51, p<0.0001) were found (**Table S6**, **Figure S3** and **Figure S4**).

**Supplementary Table S6.** Results from the meta-regressions with moderators.

| **Moderator** | **Q_M_** | **d** | **se** | **zval** | **pval** | **ci.lb** | **ci.ub** |
| --- | --- | --- | --- | --- | --- | --- | --- |
| Sample size | 0.51 | 0.00 | 0.00 | 0.66 | 0.51 | -0.00 | 0.00 |
| Mean age | 0.58 | -0.01 | 0.02 | -0.55 | 0.58 | -0.05 | 0.03 |
| Proportion of females | 0.16 | -0.68 | 0.49 | -1.40 | 0.16 | -1.64 | 0.27 |
| Proportion of first episode patients | 0.08 | 1.42 | 0.83 | 1.72 | 0.08 | -0.19 | 3.04 |
| Proportion recurrent episode patients | 0.48 | -0.64 | 0.91 | -0.71 | 0.48 | -2.43 | 1.14 |
| Proportion of AD-free patients | 0.25 | 0.64 | 0.56 | 1.15 | 0.25 | -0.45 | 1.73 |
| Proportion of AD-using patients | 0.49 | -0.45 | 0.66 | -0.69 | 0.49 | -1.74 | 0.83 |
| Proportion of remitted patients | 0.17 | -2.05 | 1.50 | 1.37 | 0.17 | -4.99 | 0.89 |
| Proportion of acute patients | 0.59 | 0.40 | 0.74 | 0.54 | 0.59 | -1.06 | 1.86 |
| FreeSurfer version 5 | **0.01** | 0.23 | 0.10 | 2.34 | **0.02** | 0.04 | 0.42 |
| version 6 |  | 0.41 | 0.16 | 2.62 | **0.01** | 0.10 | 0.71 |
| version 7 |  | -0.10 | 0.16 | -0.59 | 0.56 | -0.41 | 0.22 |
| Scanner vendor GE | **<0.001** | -0.02 | 0.16 | -0.12 | 0.91 | -0.33 | 0.29 |
| Philips |  | 0.51 | 0.12 | 4.06 | **<.0001** | 0.26 | 0.75 |
| Siemens |  | 0.10 | 0.10 | 0.97 | 0.33 | -0.10 | 0.75 |
| Field strength | 0.44 | 0.26 | 0.34 | 0.77 | 0.44 | -0.40 | 0.92 |
| R^2^ | 0.94 | 0.01 | 0.09 | 0.07 | 0.94 | -0.16 | 0.18 |
| MAE | 0.79 | 0.01 | 0.05 | 0.26 | 0.79 | -0.08 | 0.10 |

Abbreviations: Q_m_, model sum of squares; d, Cohen’s d effect size; se, standard error; zval, z-statistic; pval, p-value; ci.lb, 95% confidence interval lower boundary; ci.ub, 95% confidence interval upper boundary; AD, antidepressants; MAE, mean absolute error. Significant p-values are bold.

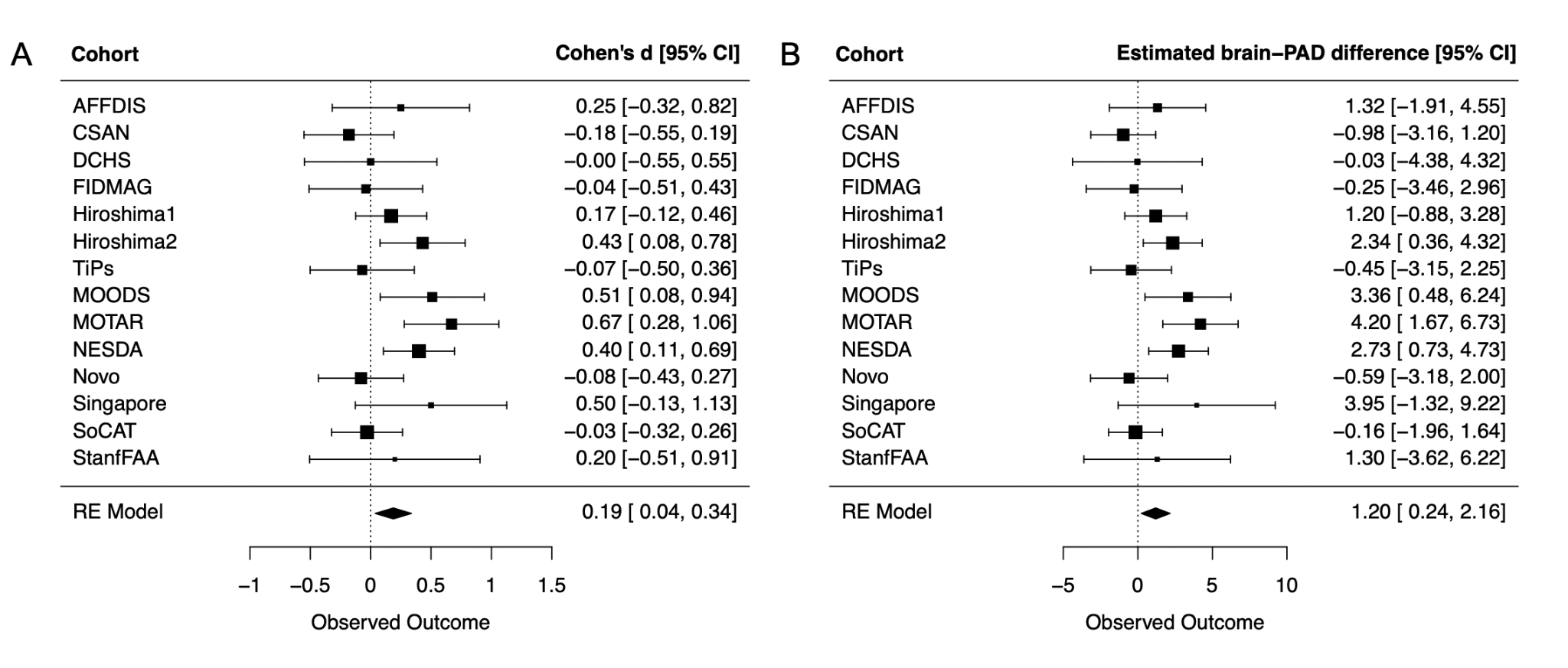

**Supplementary Figure S2.** Forest plots of case-control comparisons of brain-PAD. Panel **(A)** displays Cohen's d effect size estimates and panel **(B)** shows the estimated brain-PAD difference in years within each individual cohort. Pooled effect size (Cohen’s d=0.19, 95% confidence intervals [0.04, 0.34]) and estimated gap (b=1.20 years, 95% confidence intervals [0.24, 2.16] in depression of this meta-analytic approach is highly similar to the pooled effects from the mega-analytic approach reported in the main manuscript.

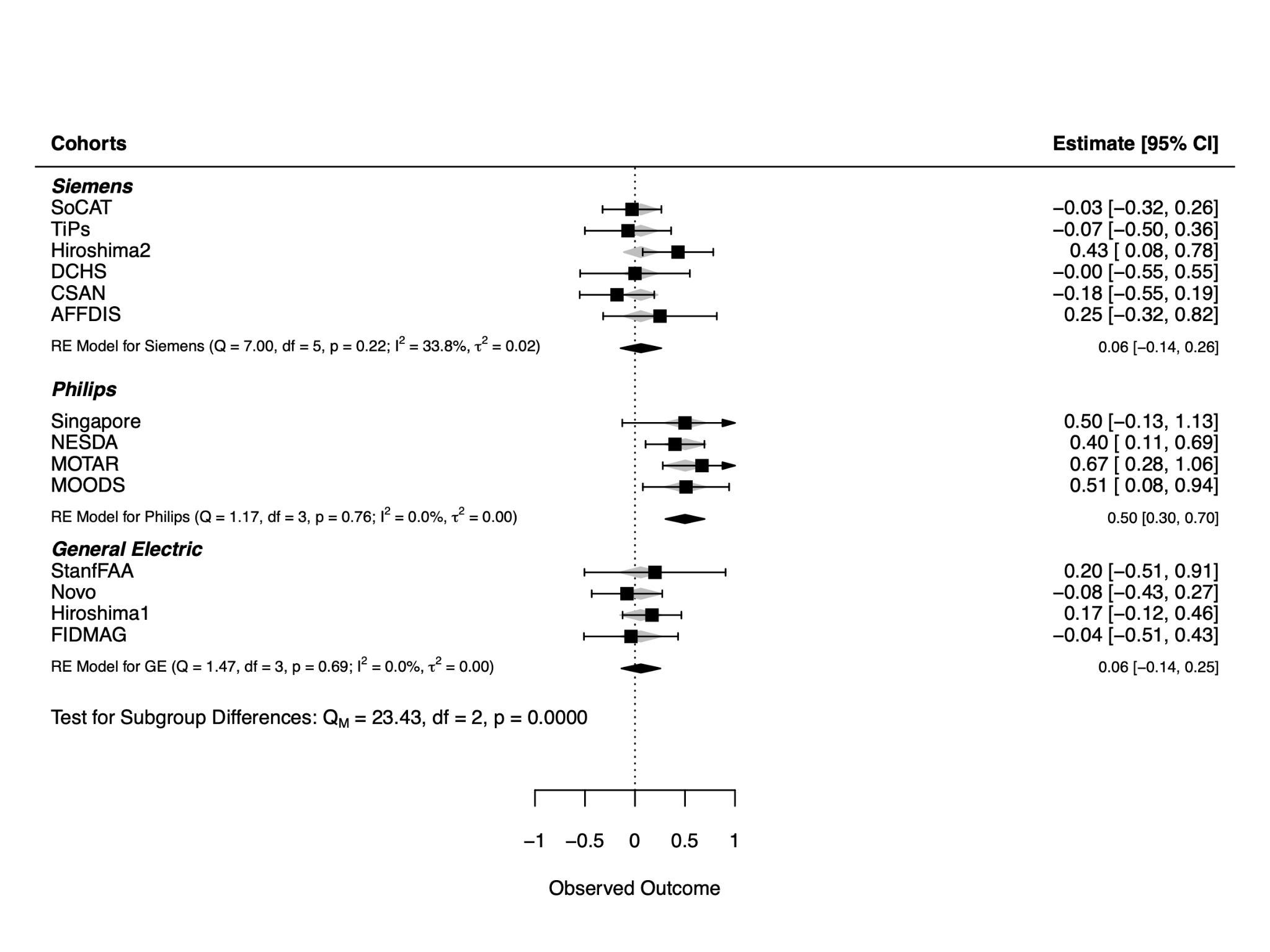

**Supplementary Figure S3.** Forest plot with three subgroups of FreeSurfer version used for preprocessing. Observed outcome and estimate indicate Cohen’s d effect size.

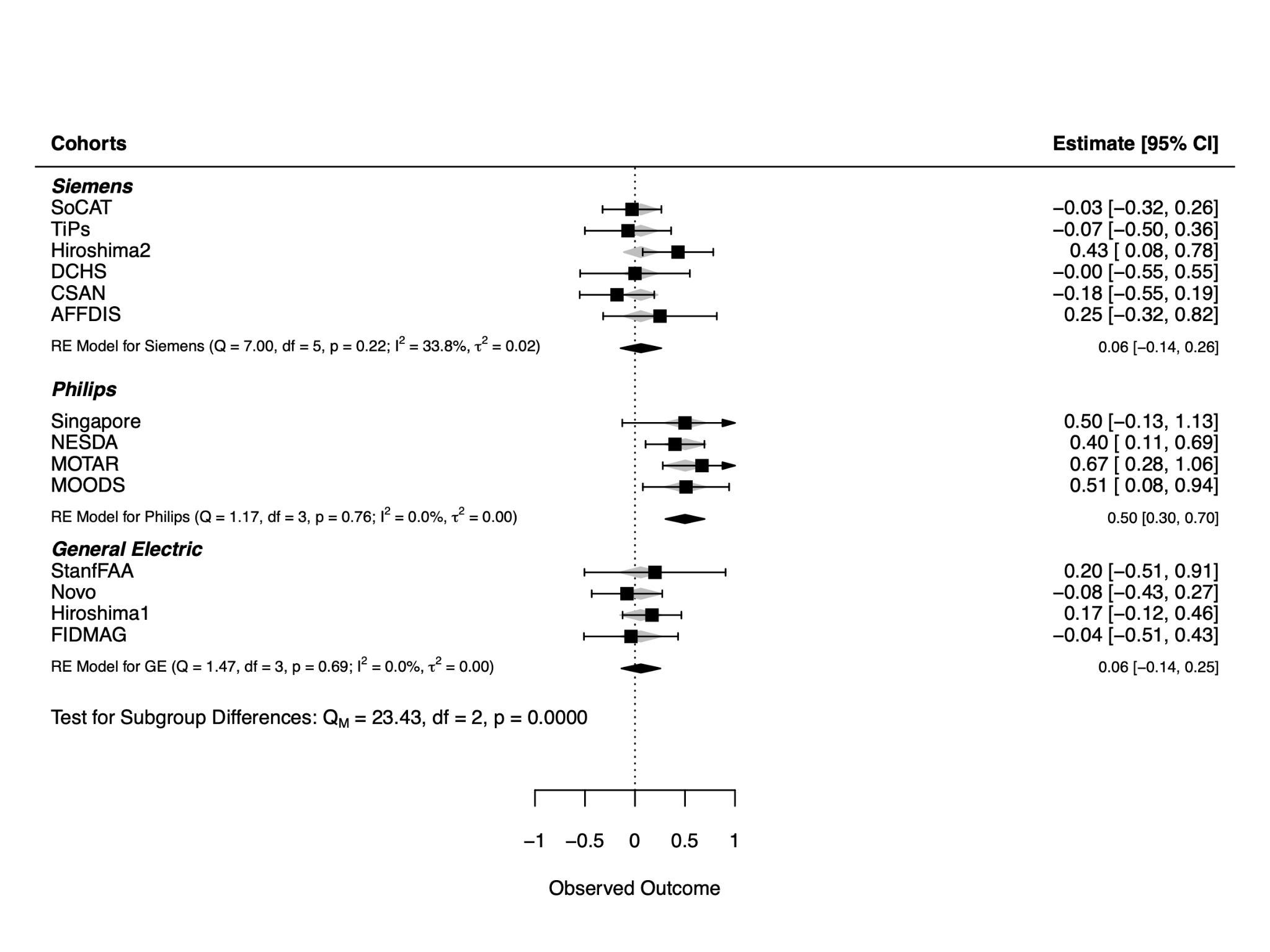
 **Supplementary Figure S4.** Forest plot with subgroups of scanner vendors used for image acquisition. Observed outcome and estimate indicate Cohen’s d effect size.
